# Is Overfishing the Main or Only Factor in Fishery Resource Decline? The Case of The Magdalena River Fishery and Its Correlation with Anthropic Pressures

**DOI:** 10.1101/2020.06.04.134072

**Authors:** Sandra Hernández-Barrero, Mauricio Valderrama Barco, Carlos Guillermo Barreto Reyes, Luis Sierra Sierra, Wolfgang Stotz

## Abstract

Overfishing has been historically considered as the main cause of fish stock depletion worldwide. This paradigm has oriented fishery management towards a classical approach, under which externalities to fisheries were not considered as they were difficult to assess and measure. The aim of this study is to describe the dynamics of different environmental, economic, and demographic variables (water flow, forest cover, gold production, population growth, stored water volume, and sediments) in relation to the behavior of the fishery production in the Magdalena-Cauca river basin from 1980 to 2015. Generalized Additive Models were used to determine the variables that best explain fishery production. The findings confirmed that environmental deterioration of the Magdalena River basin explained at least 60% of the reduction in fishery production. Thus, we concluded that the traditional approach of making fishers responsible for the decline of fish production was a misguided argument, and before implementing restrictions on fishing activity, a better understanding of the overall system is crucial. Hence, fishery management should involve the economic and social sectors that affect the offer of ecosystem services within the basin, including fishing.

## Introduction

Historically, overfishing was considered to be the main cause of fish stocks depletion in the world. This fact guided fisheries management towards a classical approach that did not adequately consider the impacts of external factors, either because they were difficult to control (1,2), or complex to characterize. This is the case in inland fisheries, being rivers the most impacted ecosystems by human activities over the past 100 years. Furthermore, many activities linked to the use of natural resources that imply intense human interventions take place in rives, threatening their functionality (flow rates disruption, erosion, alterations of habitats, among others) (3). As a consequence, it is imperative to study the anthropic effects on both, the environment as well as on natural fish populations, before ascribing all impacts to fishing activity (4). Therefore, as fisheries cannot be considered isolated, a multifactorial approach is required for their assessment and management (5).

According to the abovementioned, inaccurate or incomplete diagnoses of the root causes of overfishing can lead to errors in the formulation and implementation of fishing policies or programs (6). This phenomenon responds to a lack of knowledge on the impacts of other sectors (i.e., agriculture, mining, and transport, among others) in inland fisheries, which together with a northern hemisphere industrial fisheries approach (1) have resulted in the overall reduction of fishery resources. Moreover, these concepts are focused on internal factors such as size, fishing gear, reproductive seasons, and reserve areas, aimed to achieve sustainability only through their management (1,7).

Within this reference frame, artisanal fisheries in the Magdalena River basin show a decline in their discharges from average production of 70 000 t per year (in the 1970s) to about 30 000 t per year (8). In the same time, the basin has been altered by various activities potentially causing environmental impacts on fish populations and, therefore, on fishing activity. Moreover, according to Restrepo and Restrepo (9), 63% of the original ecosystems of the basin have been altered. Rodríguez-Becerra (10) found that 80% of the GDP of Colombia, 70% of the hydraulic energy, 95% of the thermoelectricity, 70% of the agriculture and 90% of the coffee are produced in the basin of the Magdalena River; however, the effect of all these activities on fishing production remains unknown. This development within the basin could affect fish breeding, survival, and development of their larvae and juveniles, as well as their feeding dynamics.

The breeding behavior of fish species, particularly migratory ones –which contribute to a large proportion of the fish production in the Magdalena River– is directly related to water flow. Fish react physiologically to the hydroperiod and flooding cycles of rivers (11,12), which determine the interconnectivity of the aquatic environments in which fishes perform their breeding migrations (11,13,14). But the hydroelectric power development of the country fragments the river, interrupting the reproductive migrations that in the Magdalena-Cauca basin reaches an altitude of 1,200 m above the sea level (14). Dams, storing water volumes below that altitude, affect the migrations directly and, consequently, the fishing production of those species. Likewise, another pressure factor that could be affecting fish reproduction is the presence of mercury, a gold mining subproduct, which significantly contributes to the overall contamination in the basin (15). This heavy metal is incorporated in the food chain of fishes generating impacts in their reproductive health (16,17). As a result, an inverse correlation is expected to be found between fish production of all species and variations on water flow regimes, stored water volumes, and mercury concentrations.

Fish production in a floodplain river system is also related to the environmental conditions under which fish larvae and juveniles develop. Thus, the environmental deterioration of these habitats will affect population dynamics, especially in terms of recruitment and growth. These habitats are impacted by human population growth that results in an increasing demand for the use of rivers and their surrounding land.

Moreover, demographic growth also disturbs the structure of aquatic ecosystems, diminishes their integrity, and influences the capacities of fish and other organisms to survive (18,19), affecting, in turn, the overall ecosystem functions (3). Therefore, as a result, we should find an inverse correlation between demographic growth in the basin and the abundance of fish in the river.

It is evident that the natural productivity of aquatic environments sustains fishery production, and that human intervention produces changes in the regimes and/or dynamics of that productivity; this occurs in particular by the increase of nutrient concentrations in the water as a result of changes in land use that also facilitate sediments transport. Thus, the clearcutting of forests leads to major waste-generating activities and multiplies erosion processes. Up to 79% of the catchment area of the Magdalena River suffers severe erosion conditions partly because of the deforestation of more than 70% of its natural forests, a process that took place from 1980 to 2010 (20). According to Kjelland et al. (17), high sediment loads in the water affect the feeding behavior of fish and impacts the trophic structure (predator-prey relation). Therefore, an adverse effect of decreasing forest cover on fish populations would be expected.

Accordingly, the aim of this research was to respond to how much and to what extent the external factors to fisheries affect fish production.

## Materials and Methods

### Description of the study area

The Magdalena River basin is the main watershed in Colombia (South America), with a drainage area of 257,400 km^2^, comprised by two large inter-Andean rivers, Magdalena River (the most important of its nature in South America) and the Cauca River; both have high, medium and low basins (21) (Fig 1). These water bodies are located within a broad spatial and altitudinal distribution (from 3,685 m down to sea level) and encompasses all ecosystems present in the Andean and Caribbean regions, with a varied and complex mosaic of biomes, resulting in a diversity of environments and organisms (22).

**Fig. 1.**
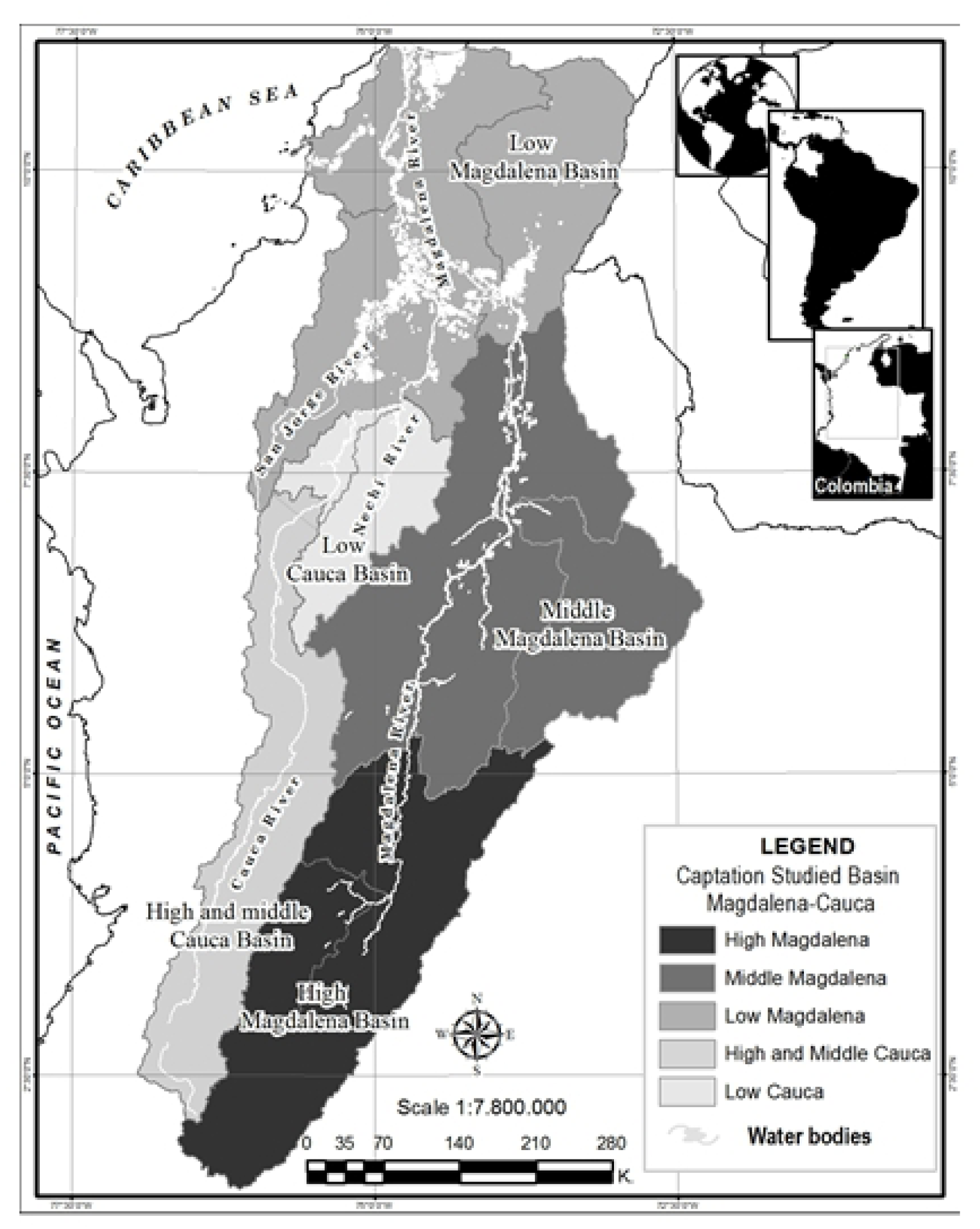
Study basin Magdalena-Cauca. We present the five hydrographic zones that cover the Magdalena-Cauca river basin.

One hundred fifty-one sub-basins comprise the basin, 42 of which are of second-order (23), with flood plain systems covering a wide variety of environments (marshes, sandbanks, lakes, lagoons, streams, reservoirs, artificial water bodies, and channels), which form interconnected ecological units (24). In this system, the fishing activity provides at present an estimated total annual production of 30,000 t, representing a commercial value in 2010 of COP 368,863 million (US$204 million) and providing food security for more than 175,000 people (24).

In terms of basin occupation, agricultural areas are predominant, representing 58.20%, followed by forest and semi-natural areas (28.71%), urban and industrial areas (0.85%), wetlands areas (2.75%) and water bodies (2.57%), being this last one of vital importance in the basin context (9,21). The basin crosses 19 departments and 128 municipalities and offers the ideal conditions for the development of several important urban centers, with a total population of 32 million people.

### Predictor environmental variables

Different environmental and demographic variables were analyzed (water flow, forest cover, gold production, demography, stored water volume, and sediments) in relation to the behavior of fishing production in the Magdalena-Cauca river basin, to study how environmental changes impacted fishing production between 1980 and 2015.

These variables were selected according to the following criteria: water flow changes and stored water area as they affect migratory patterns and natural dynamics of species; mercury, as it affects the reproductive health of species; forest cover, demographic growth, and sediments as they are considered as a proxy for changes in water quality, alteration of environments and modification of natural productivity regimes generated by the increased nutrient content in its waters.

The historical information of the annual hydrological data (1975-2015) was provided by Instituto de Hidrología, Meteorología y Estudios Ambientales (IDEAM) [Institute of Hydrology, Meteorology and Environmental Studies IDEAM] and included monthly average water flow series obtained at the Calamar weather station on the main channel in the lower Magdalena River basin. The database of the total stored water volume in the basin was compiled by our team based on the list of water reservoirs in the basin, information provided by Jiménez-Segura et al. (2011); the parameters considered were: year of construction, effective volume (M·m^3^) and altitude (meters above the sea level). These values were estimated for the area above and below 1,200 m a.s.l., as this altitude is considered as the limit for migratory fish distribution. To analyze mercury concentration associated with gold mining, we considered the total annual gold production in the department of Antioquia in kg·year^-1^ since more than 60% of the gold (vein and alluvial) produced in the Magdalena River basin comes from this region. The data was obtained from Sistema de Información Minero Energético (SIMCO) [Mining and Energy Information System (SIMCO) (https://www1.upme.gov.co/InformacionCifras/Paginas/Boletin-estadistico-de-ME.aspx)], and the historical records from Ministerio de Minas y Energía [Ministry of Mines and Energy) (https://biblioteca.minminas.gov.co/pdf/1989%20-%201990%20MEMORIA%20AL%20CONGRESO%20NACIONAL%20ANEXO%20HISTORICO.pdf)].

Regarding the forest area, this value was established according to the databases provided by IDEAM that describe the area covered by natural forest (ha·year^-1^) in the Magdalena River basin between 1990 and 2016. The historical demographic records (number of persons·year^-1^) of the 19 departments within the basin, were provided by Departamento Administrativo Nacional de Estadística (DANE) [National Administrative Department of Statistics DANE], based on five censuses with projections until 2020. Regarding sediments, we analyzed them as solid flow, which is an indicator of the material transported by the water current, and registered in the main channel of the Magdalena River in its lower basin. The annual sediment transport was estimated based on the average daily time series per month (Qs·t·d^-1^) (20,23).

### Response variable

The response variable analyzed was the total fishery production of the basin in t·year^-1^ for the period 1975 to 2015, presented by Barreto (8). This database allowed us to estimate the percentage contribution of migratory and non-migratory species to fishery production over time.

### Information processing and analysis

The variables were tabulated in a single database. The starting point for the analysis was the year 1980, and we included variable entries until the year 2015. The gaps in the information were filled with synthetic data estimated using the regression trend technique. In all cases, both retrospective and prospective projections showed an error within a range of about 1%.

With the collected and estimated information, we built a 36 x 9 matrix (years x variables) (S1 Text). We first scanned the data to establish the distribution pattern of the variables (with or without normal distribution). Datamining established non-parametric models as a path, so we decided to use the Generalized Additive Models (GAM) of Hastie and Tibshirani (25). This technique allowed the adjustment of statistical models, as well as the study of natural phenomena with non-linear complexity behavior (26), which, aligned with the ecological theory (27), provides a greater possibility of controlling the confounding variables (28,29). GAM corresponded to the following equation:

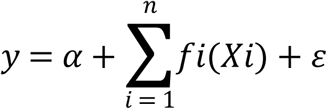

Where *y* was the response variable, *Xi* the predictors, α a constant and ε the error. The *f*i parameters for the non-parametric and Gaussian functions were estimated using smoothing spline (*s*) based on (27). In the GAM diagnostic process, we included the significance value (*p*), the calculation of the percentage of deviance explained by the model, and the Akaike information criterion (AIC). The software used was R (30). Thus, with the response variable (fish production) and the different predictor variables, we created 203 combinations allowing us to run 35 models (S1 File). We selected the model that best explained fish production behavior in the Magdalena River basin, based on the deviance and the AIC values. For precision purposes and considering the variable stored water volume, the five models that simultaneously crossed “water volume stored below 1200 m a.s.l., and total stored water volume for the entire basin” were discarded, as the second includes the first.

Considering how relevant water flows are for fish, and because flow patterns have a bimodal behavior in this particular basin, we estimated the maximum water flow value for each semester of every year and then calculated the difference among those values for the period between 1975 and 2015. We graphed the water flow differences vs. fish production and applied a two-period moving average.

## Results

Fishery production in the Magdalena River showed two periods of abundance; the first period (1980-1991) with higher fish production records but larger fluctuations, and the second period (1992-2015) with lower fish production records and smaller fluctuations with a trend towards stability (Fig 2). According to the statistical records, non-migratory species began to be reported in the early 1990s, showing an average contribution to fisheries production of 4% compared to the 96% for migratory species. Non-migratory species represent 11% of the fisheries production between 2010 and 2015.

**Fig. 2.**
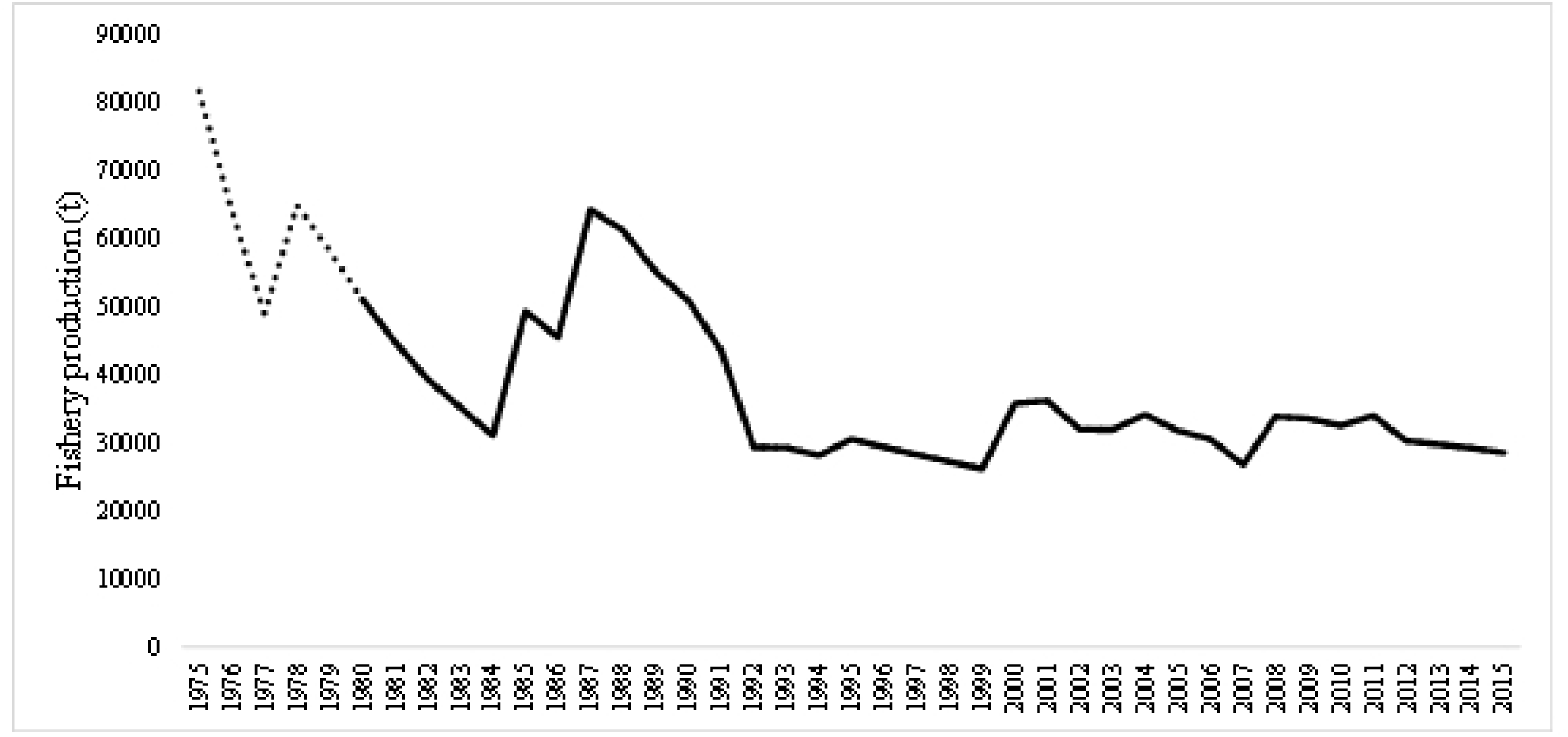
Estimated fishing production (t) for the Magdalena-Cauca River Basin between the years 1975 and 2015. The data analyzed corresponds to the period from 1980 to 2015 (dark line). Source: (8).

The correlation between fisheries production and the differences in the maximum amplitude of water flows showed a coupling between both variables (Fig 3). In general terms, the fishery production curve presented the same shape and pattern as the curve generated by the moving averages; that result suggests a strong influence of water flow regime on fish abundance in the system (Fig 3). In 1992, after significant fluctuations, the average water flow decreased by 1% during the dry season and increased by 7% during the rainy season. Regarding the magnitude of the water flow differences observed between the years, after 1992, the magnitude of the differences became more stable and coincided with a period of a low and stable fish production (Fig 2 and Fig 4a).

**Fig. 3.**
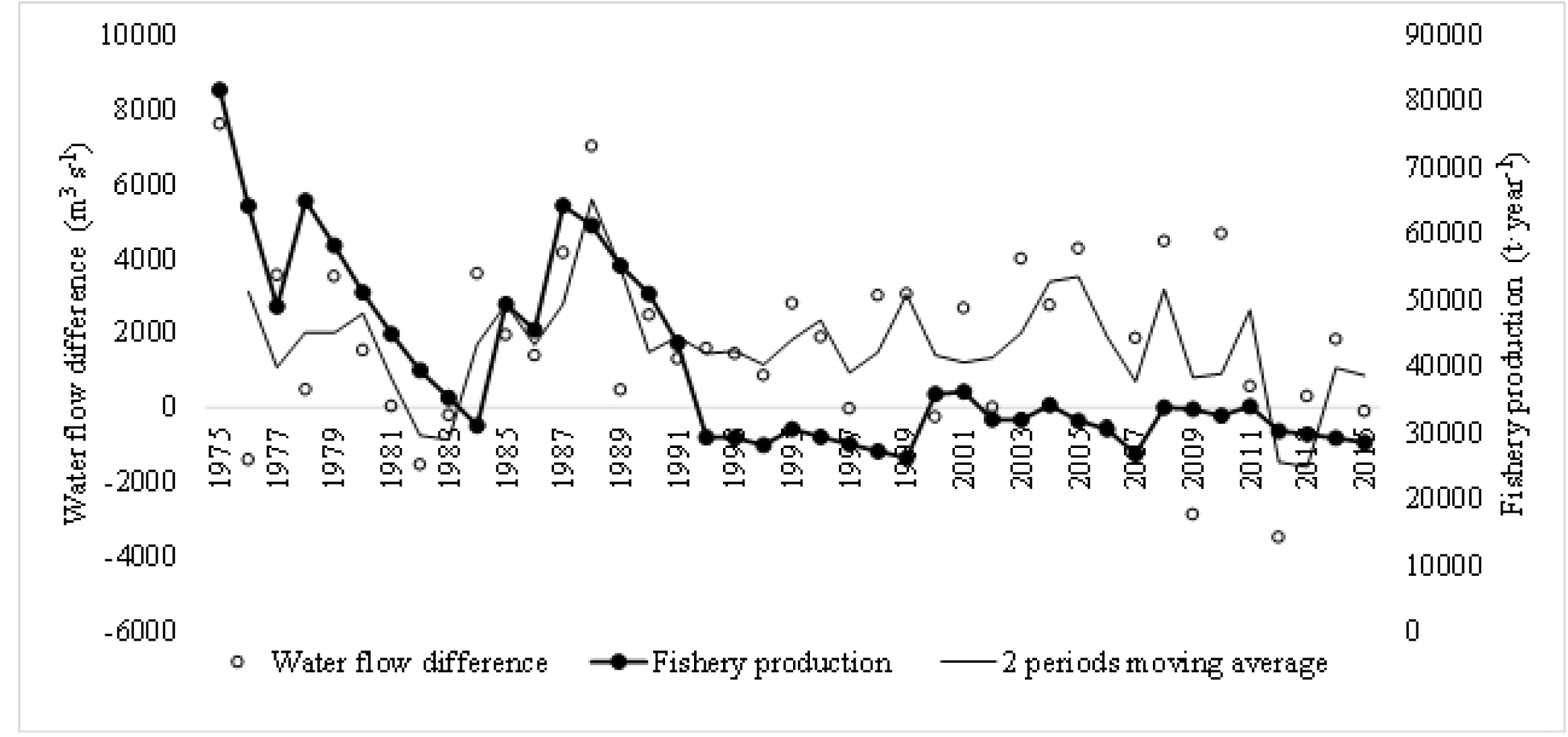
Moving average between the difference of maximum water flow rates in the first and second semesters of each year and its correlation with fishing production between the years 1975 and 2015. The thick black line shows the behavior of fish production over time and the thin black line the maximum water flow value for each semester of every year.

**Fig. 4.**
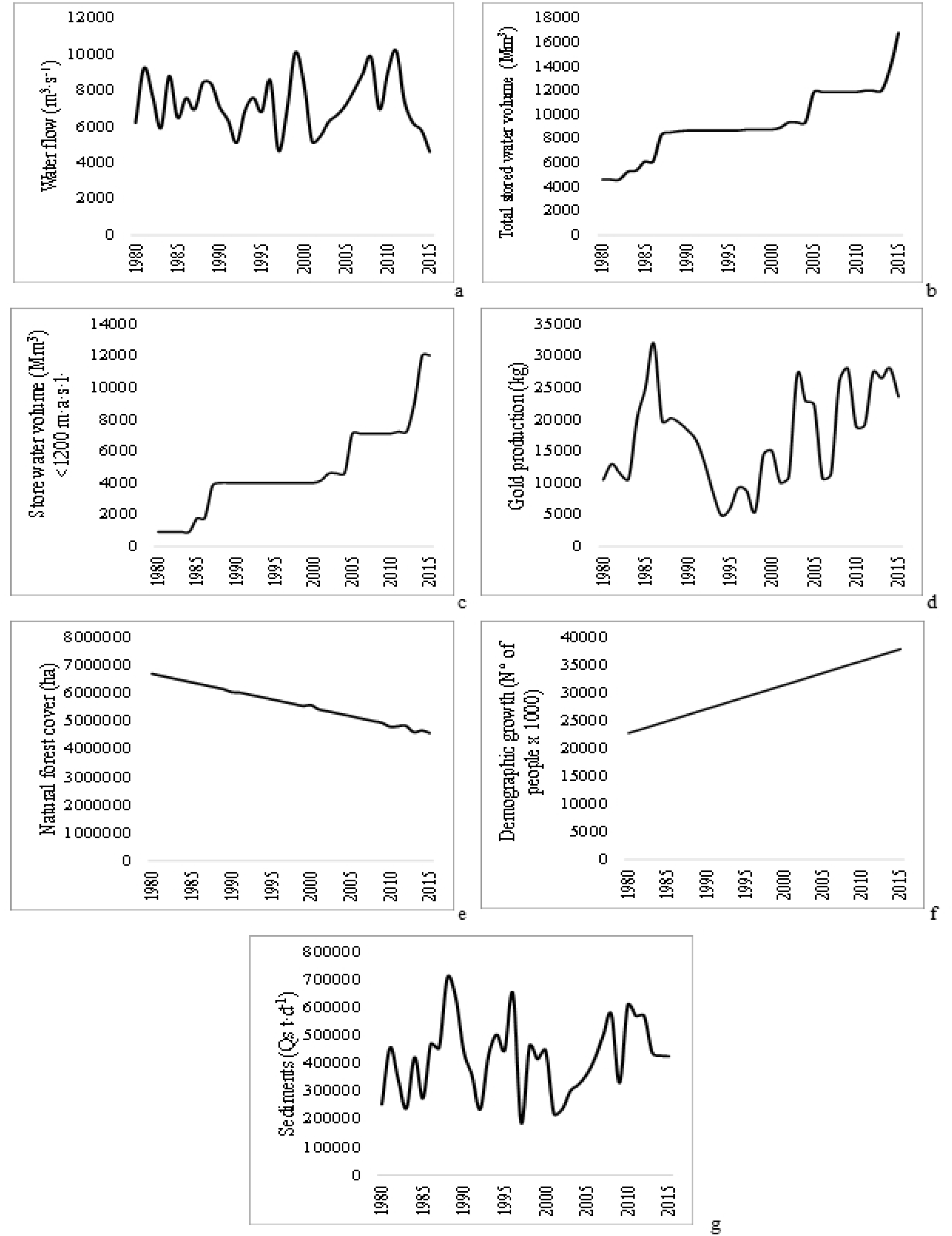
Behavior of predictor variables during the period between 1980 to 2015 in the Magdalena-Cauca River Basin. (a) water flows (m3·s-1); (b) total stored water volume (Mm3); (c) stored water volume < 1,200 m a.s.l; (d) gold production (kg); (e) forest cover (ha); (f) demographic growth (number of people x1000); (g) sediments (Qs·t·d-1).

It is noteworthy that the period when the difference in water flow became minimal (from 1982 to 1983), was followed by a significant reduction in fish production (Fig 3); likewise, in the following years, when the difference in water flow recovered, the same happened with the fish production. From 1992 and onwards, when reductions and fluctuations in water flow differences decreased and remained steady, fisheries production also behaved in this same way.

Regarding the volumes of stored water, for both the total and below 1,200 m.a.s.l., these values showed a gradual increase with strong increments in the periods 1986-1988 and 2004-2006; eventually, the most significant increase occurred from 2013 to the present day (Fig 4b and c). These increases are related to the progressive implementation and operation of the main dams in the basin. In turn, the first main increase in the total water volume stored and the one below 1,200 m a.s.l. occurred in 1986 and concurs with the reduction in fish production, suggesting a change in the environmental dynamics of the river that prevented fish populations from recovering their previous abundance (Fig 2 and 4c).

Gold production showed a rapid increase between 1980 and 1986, followed by a steady decrease until 1994, before showing a continuous but fluctuating upward trend (Fig 4d). It should be noted that fish production decreased right after gold production reached its highest value (Fig 2), suggesting a slightly delayed effect of mercury contamination due to gold mining on fish stocks.

The reduction of the forest area was continuous and progressive during the entire study period (Fig 4e). The same can be observed for demographic growth (Fig 4f).

Concerning the sediment fluctuation over time, the change in the magnitude of the maximum values after 1986 is noteworthy (Fig 4g). From that year onwards, the maximum values are approximately 30% higher than in previous years, corresponding also with the moment when the first dam started operations and with the decrease in fishing production (Fig 2).

Figure 5 presents the functional correlations between fish production and the predictor variables. Of the 30 models that were selected, 27 reported synergy between the different variables and presented an explained deviance of more than 60% on fish production behavior (Table 1).

**Table 1.**
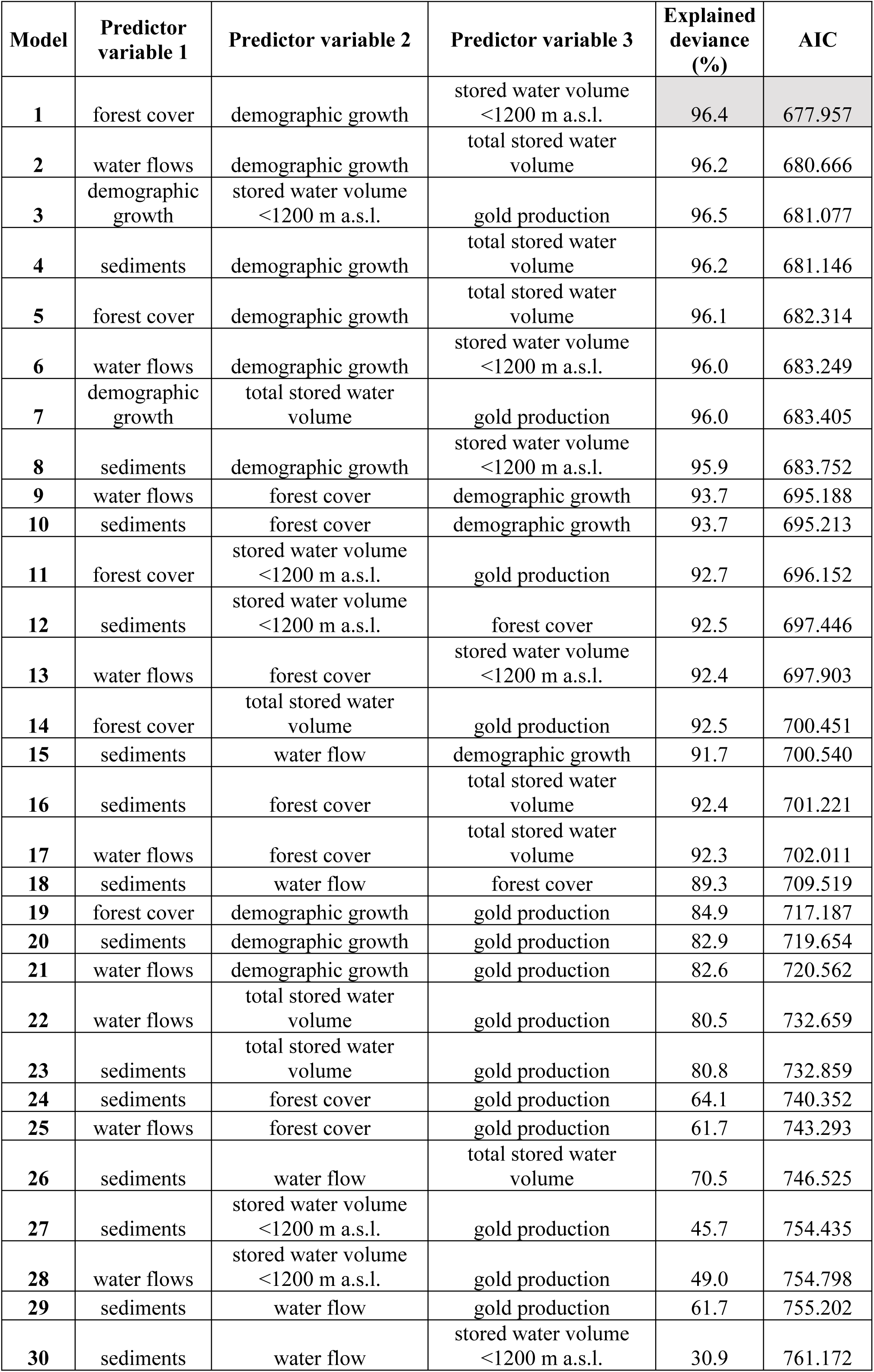
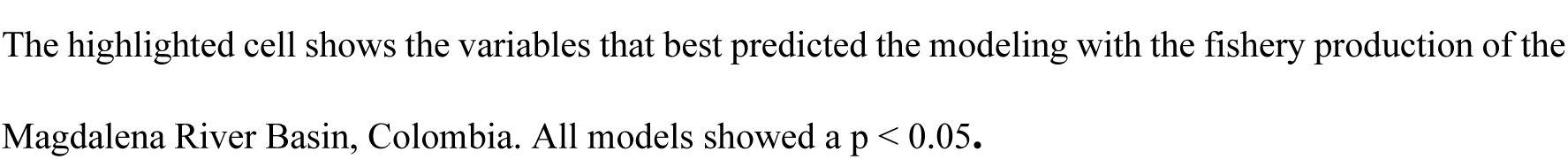
Results of the Generalized Additive Models (GAM) modeling between fishery production (t) and predictor variables, the percentage of the explained deviance by the model, and the AIC.

| Model | Predictor variable 1 | Predictor variable 2 | Predictor variable 3 | Explained deviance (%) | AIC |
| --- | --- | --- | --- | --- | --- |
| 1 | forest cover | demographic growth | stored water volume <1200 m a.s.l. | 96.4 | 677.957 |
| 2 | water flows | demographic growth | total stored water volume | 96.2 | 680.666 |
| 3 | demographic growth | stored water volume <1200 m a.s.l. | gold production | 96.5 | 681.077 |
| 4 | sediments | demographic growth | total stored water volume | 96.2 | 681.146 |
| 5 | forest cover | demographic growth | total stored water volume | 96.1 | 682.314 |
| 6 | water flows | demographic growth | stored water volume <1200 m a.s.l. | 96.0 | 683.249 |
| 7 | demographic growth | total stored water volume | gold production | 96.0 | 683.405 |
| 8 | sediments | demographic growth | stored water volume <1200 m a.s.l. | 95.9 | 683.752 |
| 9 | water flows | forest cover | demographic growth | 93.7 | 695.188 |
| 10 | sediments | forest cover | demographic growth | 93.7 | 695.213 |
| 11 | forest cover | stored water volume <1200 m a.s.l. | gold production | 92.7 | 696.152 |
| 12 | sediments | stored water volume <1200 m a.s.l. | forest cover | 92.5 | 697.446 |
| 13 | water flows | forest cover | stored water volume <1200 m a.s.l. | 92.4 | 697.903 |
| 14 | forest cover | total stored water volume | gold production | 92.5 | 700.451 |
| 15 | sediments | water flow | demographic growth | 91.7 | 700.540 |
| 16 | sediments | forest cover | total stored water volume | 92.4 | 701.221 |
| 17 | water flows | forest cover | total stored water volume | 92.3 | 702.011 |
| 18 | sediments | water flow | forest cover | 89.3 | 709.519 |
| 19 | forest cover | demographic growth | gold production | 84.9 | 717.187 |
| 20 | sediments | demographic growth | gold production | 82.9 | 719.654 |
| 21 | water flows | demographic growth | gold production | 82.6 | 720.562 |
| 22 | water flows | total stored water volume | gold production | 80.5 | 732.659 |
| 23 | sediments | total stored water volume | gold production | 80.8 | 732.859 |
| 24 | sediments | forest cover | gold production | 64.1 | 740.352 |
| 25 | water flows | forest cover | gold production | 61.7 | 743.293 |
| 26 | sediments | water flow | total stored water volume | 70.5 | 746.525 |
| 27 | sediments | stored water volume <1200 m a.s.l. | gold production | 45.7 | 754.435 |
| 28 | water flows | stored water volume <1200 m a.s.l. | gold production | 49.0 | 754.798 |
| 29 | sediments | water flow | gold production | 61.7 | 755.202 |
| 30 | sediments | water flow | stored water volume <1200 m a.s.l. | 30.9 | 761.172 |

The highlighted cell shows the variables that best predicted the modeling with the fishery production of the Magdalena River Basin, Colombia. All models showed a $p < 0.05$ .

**Figure 5.**
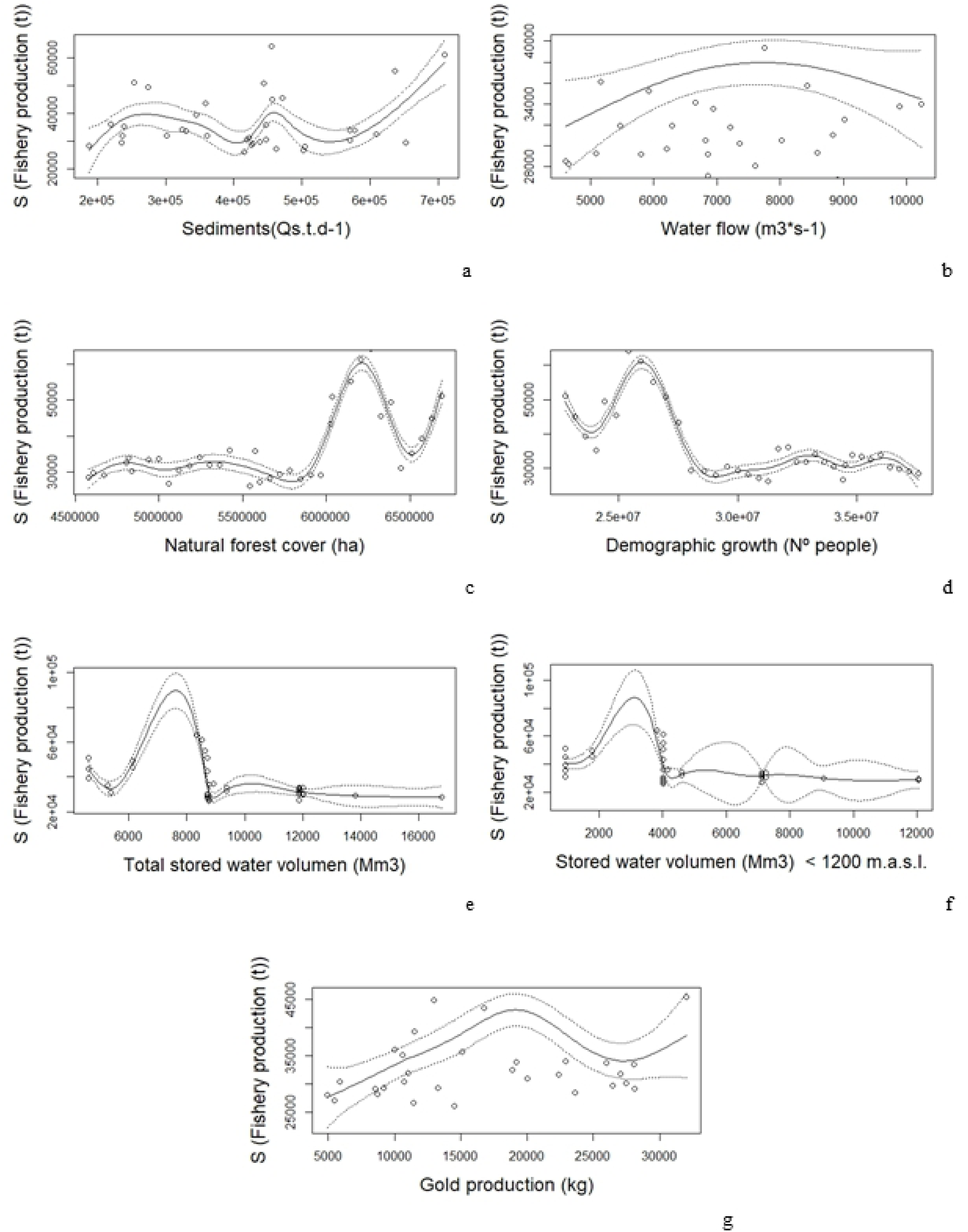
Functional correlations between fish production and predictor variables. Sediments (a), Water flow (b), Natural forest cover (c), Demographic growth (d), Total stored water volume (e), Stored water volume below 1200 m.a.s.l. (f) and gold production (g). The dotted lines represent the confidence intervals.

The environmental variable that best forecasted the fishery production was forest cover, meanwhile at a productive level, it was the total volume of stored water, and, at a demographic level, it was population growth (p < 0.05, higher explained deviance and lower AIC) (Table 2). Fishing production was high since there was a large forest cover, low total water stored volume levels, and a low population in the basin. The highest production records, which occurred between 1980 and 2015, were associated with water flows between 7,000 and 8000 m3·s-1, population densities under 26 million people in the basin, forest cover over 6 million hectares, stored water volumes of 3,000 Mm3 in reservoirs located below 1,200 m a.s.l.., total water stored volume of 7,500 Mm3, a maximum of 20,000 kg of gold, and sediment rates higher than 60,000 Qs·t·d-1 (Figure 5).

**Table 2.** GAM modeling results between fishery production and each of the type of variable.

| Type of variable | Factor | Explained deviance (%) | AIC | p-value |
| --- | --- | --- | --- | --- |
| **Environmental** | forest cover | 89.8 | 703.764 | 0.000 |
| **Environmental** | sediments | 38.5 | 765.161 | 0.100 |
| **Environmental** | water flows | 7.47 | 769.071 | 0.431 |
| **Demographic** | population growth | 88.6 | 707.182 | 0.000 |
| **Productive** | gold production | 27.9 | 764.569 | 0.100 |
| **Productive** | total stored water volume | 70.5 | 741.004 | 0.000 |
| **Productive** | stored water volume <1200 m a.s.l. | 38.5 | 766.312 | 0.110 |

Percentage of the explained deviance by the model, the Akaike criterion (AIC), and the p-value.

## Discussion

The findings confirmed that the environmental variables of the Magdalena River basin explained at least 60% of the reduction in fish production. This showed that the conventional fishery management paradigm, in which overfishing is considered to be the principal factor in the deterioration of fish stocks, is erroneous.

The most significant productive interventions were: i) the increase of human population within the basin is translated into deforestation and transformation of land use, and in turn, this alters sediment dynamics and affects the quality of the river environments and the flood plains; ii) the construction of reservoirs and dams which interfered in various ways with the reproduction processes of the main species, by modifying water flows and altering the migratory patterns, fragmenting populations with the presence of barriers, reducing dispersion and reproduction areas and, attenuating water flow regimes necessary to guarantee appropriate flood pulses for fish breeding; iii) the substantial increase in gold production which, apparently in its initial phase, generated a hazardous level of pollution affecting fish reproduction processes, and fishing production.

The variability of fish production since 1992 stabilized at a low level without recovering. The above indicates the crossing of some thresholds that not only prevented larger fish production from being sustainable but also condemning it to a negative trend. Some of these thresholds are: i) fluctuation of water flows between years exceeded 8,000 m^3^·s^-1^, ii) the stored water volume was over 3,000 Mm^3^, iii) gold production exceeded 20,000 kg, iv) the forest area decreased by 6 million ha, and v) suspended solids exceeded 400,000 Qs·t·d^-1^ at a certain point during the year.

We consider that since1992, the decrease in water flow during the upriver or upstream migration (*subienda* in Spanish) during the summer season (6.4% on average) affected the migratory and reproductive processes of the species and, as mentioned by Ellis et al. (2016), as the biological rhythms of fish are modified, the opportunities for spawning, growing and dispersing are affected. Two processes are involved in this context: water flow must decrease sufficiently for shoals to be able to swim upstream, but afterward, the water flow must increase enough to generate flooded zones where larvae and juveniles are bred (14). The abovementioned led us to highlight those “sacred” flood pulses in the Magdalena-Cauca river basin as crucial. In addition, we evidenced an indirect correlation between fish production and water flows; the modeling results showed an optimal water flow ranges (7,000-8,000 m·s^-1^), during which higher fish productions were generated. Extreme flow rates, especially for low values, are related to low productions.

Hydroelectric power plants have been built both in the main channel as well as in the tributaries of the Magdalena-Cauca rivers, altering water flows by regulating them seasonally and “daily” (pulses known as “hydropeaking”). These changes, according to Gillson (31), are agents that modify the richness and diversity of species, and therefore, fishing production in the rivers. There is a total of 39 reservoirs in the basin, storing about 16,800 Mm^3^, i.e., 73% more than the total volume stored in 1980. Thus, as the total stored water volume in the basin increased, fishing production decreased. Furthermore, in several sectors downstream of the dams, a 58% reduction of migratory species was detected in the Magdalena-Cauca Basin between 1980 and 2015. This pattern was also reported by Agostinho et al. (32) and Lacerda et al. (33) for other basins.

Considering this context, we propose that the habitat loss, the blockage of migratory routes and the loss of connectivity associated with the fragmentation of corridors between flood plains, have altered and are still altering the reproductive habitat as well as the fish recruitment and, consequently, fishing production. This is aligned with what has also been discussed and published by various authors (14,34–38) for similar systems.

When evaluating gold production as a proxy for mercury pollution, we consider that the intensive extraction that took place in the late 80s, led to a prolonged impact (contamination), as the accumulation in the sediments and bioaccumulation in the food chain continued to affect fish populations in later periods. Recent studies showed that in Colombia, between 80 and 100 t of mercury are released annually; its presence has been reported in more than 13 fish species in the basin, with concentrations higher than 0.2 μg of Hg per gram of fresh mass, making them unsuitable for human consumption (39). Mercury concentrations are generating impacts on the reproductive health of fish, detected on the alteration of sex ratios, and the reduction of the survival capacity of the offspring, also mentioned by Kjelland et al. (17) and Crump and Trudeau (16). We consider that mercury concentration is a highly hazardous factor, given its impact on human health.

Deforestation associated with the transformation of aquatic ecosystems through land-use changes has also influenced fisheries production. In the Magdalena River Basin, forest cover decreased by 32% and fishery production by 44% between 1980 and 2015. Therefore, to reach higher fish production rates, the forest area must equal what was found 28 years ago, i.e., 6.6 million hectares. Clearly, deforestation has led to changes in aquatic ecosystem conditions, which have affected fish production. This agrees with Castello et al. (40), who found that deforestation of the floodplain in the lower Amazon River basin was the main variable that explained variability in the reduction of fishing yields.

In terms of demographic growth, we confirmed the hypothesis that the higher the population growth, the lower the fish production; therefore, it can be used as a proxy for the alteration of water quality in the basin. This is because urbanization and human activities lead to large urban wastewater discharges into the rivers, which affect the water quality and life of aquatic organisms. Fishing production in the basin had its largest records in the 80s when less than 26 million people were living in the basin; however, currently, the basin has about 6 million more inhabitants. The anthropic pressure on water use (agricultural and livestock sectors), together with moderate to low water regulation, have led 66% of the area of the Magdalena River Basin to show critical, very high and high levels, as well as to a moderate to low water regulation, according to the National Water Evaluation carried out by Instituto de Hidrología, Meteorología y Estudios Ambientales (41).

According to the abovementioned and also suggested by Couceiro et al. (42) and Amisah and Cowx (18) for other regions, population growth is mirrored by the demand for river uses, which disrupts the structure of aquatic ecosystems by diminishing their integrity and influencing the ability of fish and other organisms to survive; likewise, as a consequence of this water use, eutrophication processes have been registered in the flood plains of the Magdalena River Basin such as the Zapatosa marsh (43).

Regarding sediment loads, our results indicated that fish production increased at higher sediment values. This led us to reconsider the hypothesis that sediment loads over the past two decades were stimulating accelerated sedimentation processes in the marshes. This situation is observed in the field and reported by the fishers as one of their biggest challenges. Nonetheless, Restrepo et al. (20) indicate that despite increased erosion, some of the sediments are being retained in the tributaries and, therefore, never reached the main channel (Magdalena River) as previously assumed. However, there is no doubt that sediments are related to flood pulses and nutrient cycles to the extent that there is a correlation between these sediments, flood pulses, and fish production.

At the same time, Baran et al. (5), in other systems such as the Mekong River, demonstrated that a reduction of 80% of the sediment input decreases total fish biomass by 36%. Historical records in the Lower Magdalena River Basin indicated that in the years after 1993, sediment input decreased in both the dry and wet months. Further, Jiménez-Segura et al. (14) warn us about changes in the sediment dynamics of the river due to the future implementation of hydroelectric generation projects (205 new dams by the year 2027) that will increase the energy production in Colombia by a factor of four (24 000 MW).

This reality in the Magdalena-Cauca river basin has encouraged urgent appeals to Colombian environmental institutions to implement real strategies for fisheries management with an ecosystem, inter-sectoral (44) and fish conservation approach, considering that the trans-Andean basins of the Caribbean region are the core of the economic development of Colombian society (45). We agree that a basin-wide approach is necessary, including cumulative effects and also climate variability, which merits immediate and coordinated intervention within the framework of strengthening inter-sectoral governance of fisheries. It is clear that the decline in ecosystem services and the associated severe socio-economic and environmental impacts will be increasingly challenging to reverse or mitigate these, affecting thousands of coastal inhabitants whose livelihoods depend or not on fisheries.

To summarize, the results obtained allowed us to conclude that the decrease in fish abundance was in large proportion due to environmental causes. We consider that fishing activity and landings responded more to the environmental state of the ecosystems than to any sort of approach in fisheries management. We consider that fishers in recent years have self-regulated towards new levels of abundance and that fishery authorities should be more supportive towards good fishing practices that the fishers have adopted for their survival, as a result of the reality they perceive every day. Surely, making fishers responsible for the decrease in fish production is a misguided argument, and, before trying to implement restrictions on fishing activity, a better understanding of the entire system and its dynamics is necessary. Recently, different approaches have been discussed, like the concept of balanced exploitation (46), which considers that fishing pressure should be distributed in proportion to the natural productivity of ecosystems, forcing, in our case, a response to environmental dilemmas. We consider that the system has already been adjusted to a lower level and found a new balance. Therefore, the implementation of classical fisheries management based on the overfishing paradigm is no longer sustainable, and managers of artisanal fisheries can no longer avoid external factors. Moreover, fisheries management must involve the different economic and social sectors that affect the different ecosystem services provided by the basin.

## Supporting information

**S1 Text Database of the different environmental, productive, and demographic variables analyzed concerning the behavior of fishing production in the Magdalena-Cauca river basin**. The total fishery production of the basin in t·year^-1^ was obtained of (8). The average water flow series were provided by Instituto de Hidrología, Meteorología y Estudios Ambientales (IDEAM) [Institute of Hydrology, Meteorology and Environmental Studies (IDEAM)]. The total stored water volume in the basin (effective volume M·m^3^) was compiled by our team based on the list of water reservoirs in the basin, information provided by (47). These values were estimated for the area above and below 1,200 m a.s.l.. The total annual gold in kg·year^-1^ was obtained from Sistema de Información Minero Energético (SIMCO) [Mining and Energy Information System (SIMCO) (https://www1.upme.gov.co/InformacionCifras/Paginas/Boletin-estadistico-de-ME.aspx)], and the historical records from Ministerio de Minas y Energía [Ministry of Mines and Energy) (https://biblioteca.minminas.gov.co/pdf/1989%20-%201990%20MEMORIA%20AL%20CONGRESO%20NACIONAL%20ANEXO%20HISTORICO.pdf)]. The forest area (ha·year^-1^) between 1990 and 2016 was provided by IDEAM. Between 1980 to 1989 the gaps in the information were filled with synthetic data estimated using the regression trend technique. Prospective projections showed an error within a range of about 1%. The historical demographic records (number of persons·year^-1^) of the 19 departments within the basin, were provided by Departamento Administrativo Nacional de Estadística (DANE) [National Administrative Department of Statistics DANE]. The annual sediment transport (solid flow) was estimated based on the average daily time series per month (Qs·t·d^-1^) (20,23). (DOCX)

DOI 10.17605 / OSF.IO / 268XB

**S1 File. Contains scripts used to develop the generalized additive models (GAMs)**. The application of the GAMs also included the development of various tests such as verification of residual deviance against the theoretical quartiles, analysis of residues against the line of prediction, histogram of the residues, graph of the response variable against the estimated values. Likewise, we checked the handling of multidimensional planes (K, edf, k-index and p-value). The software used was RStudio (FILE R) DOI 10.17605 / OSF.IO / 268XB

## Author Contributions

Conceptualization: SHB MVB LSS WS, Methodology: SHB CGBR, Formal analysis and Data Curation: CGBR, Investigation: SHB, Writing - Original Draft: SHB LSS, Project administration: SHB, Writing - Review & Editing: MVB WS, Supervision: WS

## Acknowledgements

We thank Fundación Humedales for supporting the main author and her technical team in the field research and surveys. To Dr. Darío Restrepo for sharing with us and let us use his sediment database. To Dr. Katty Camacho for reviewing the paper and Julia Pérez Sillero and Karen Amaya Vecth for language support. To the Instituto de Hidrología, Meteorología y Estudios Ambientales - IDEAM-(Institute of Hydrology, Meteorology and Environmental Studies) for the provision of environmental information variables of the Magdalena - Cauca river basin. To the UNED-Costa Rica DOCINADE doctoral program for its guidance to the main author

